## Supporting Figures and Tables for "Molecular QTL are enriched for structural variants in a cattle long-read cohort": Supplementary information.pdf

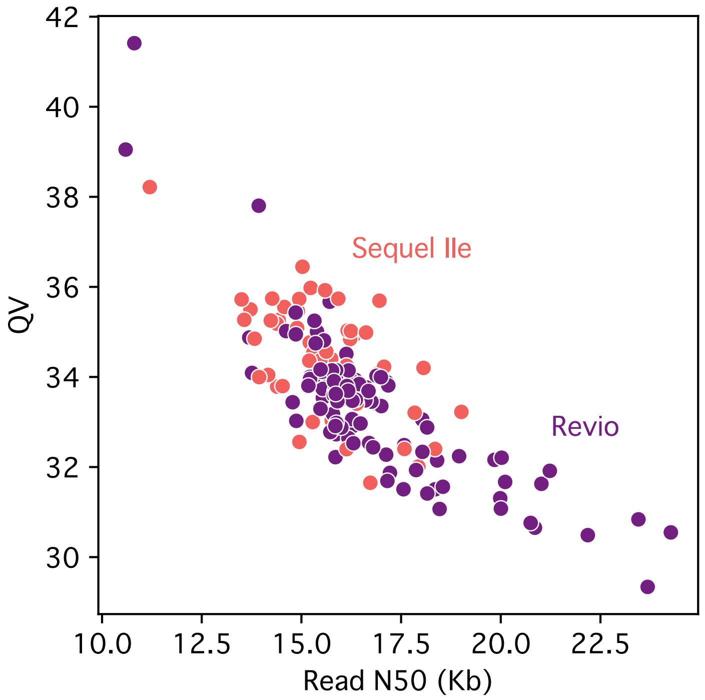


*Supplementary Figure 1.The expected negative correlation (Pearson's r=-0.80, p=5.87e-37) between read length and quality values (QV) was observed. No bias was detected across the two PacBio sequencing platforms used, with the longer read N50s for some Revio samples due to differences in library preparation rather than sequencing platform.*

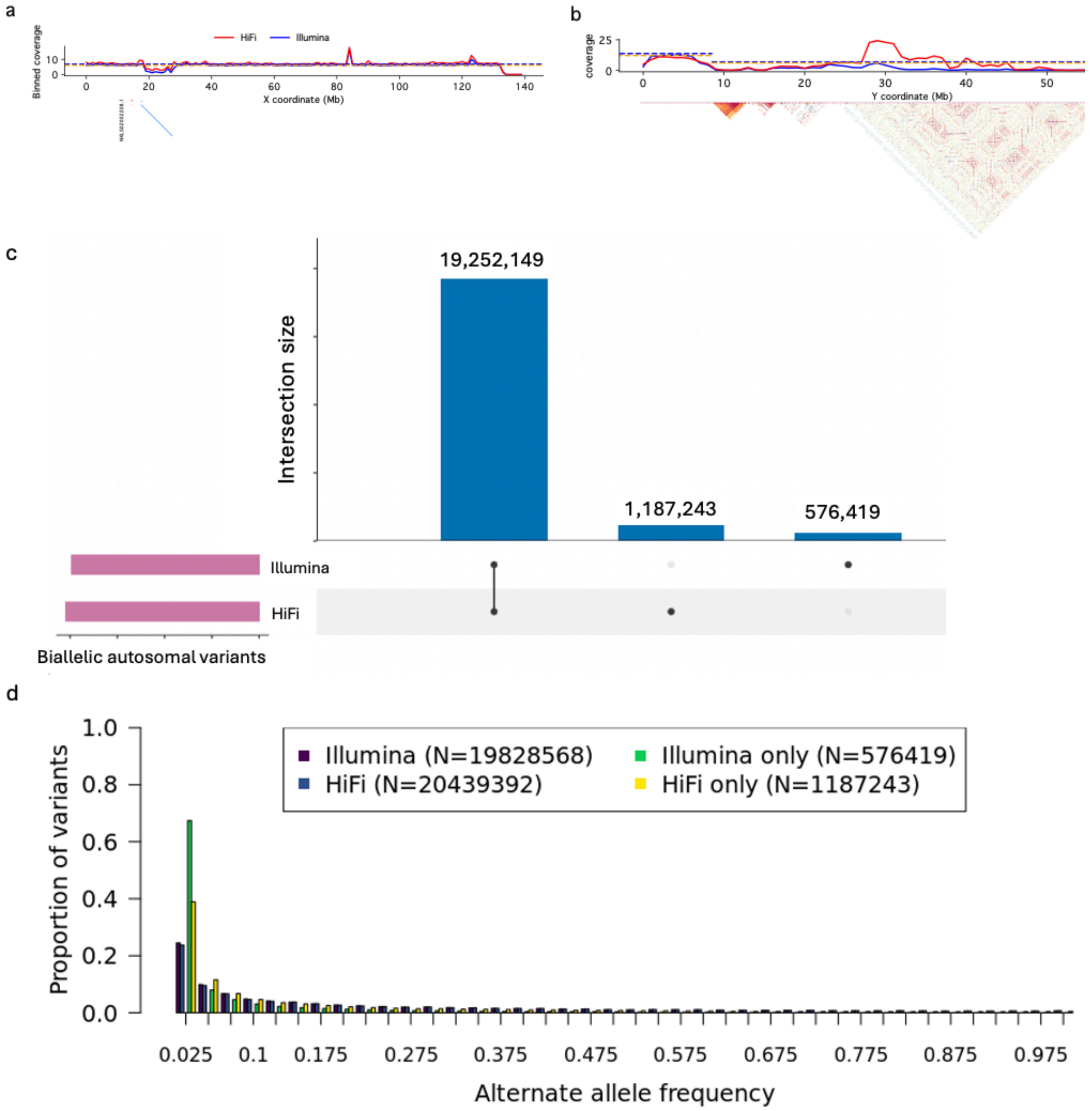


*Supplementary Figure 2. Alignment and variant calling from HiFi and Illumina reads.(**a**) Alignments to the X chromosome dipped in coverage around 17-28 Mb due to high similarity (>98%) to almost the entire length of the unplaced contig NKLS02002208.1 (11.7 Mb of the 13.9 Mb contig length). Coverage drops at the end of the X chromosome due to hard masking the X pseudo-autosomal region (X-PAR). The dashed lines indicate the expected average coverage for HiFi and Illumina reads. (**b**) In the Y pseudo-autosomal region (Y-PAR; 0-6.82 Mb), the expected coverage is higher and contains few self-similar sequences. Regions in the male-specific Y (MSY) with dense self-similar repeats, shown by dark red triangles, have low coverage. HiFi reads are better able to align to some MSY regions, although the coverage is still uneven. c) Variant calling with DeepVariant identified 21,015,811 biallelic autosomal variants in the HiFi and Illumina alignments of the 120 samples, of which 91.61% (n=19,252,149) were common to both sequencing technologies. The number of private variants was twice as high in the HiFi (n=1,187,243) than Illumina set (n=576,419). c) Of the 19.25 M common variants, 8.44% and 8.32% were detected as singletons with Illumina and HiFi, respectively (i.e., they were identified in the heterozygous state in only one individual). More than half (53.94%, n=310,937) of the Illumina-only variants were singletons, whereas the HiFi-only variants contained only 18.19% (n=215,924) singletons. This pattern suggests a higher error rate in the variants called from Illumina.*

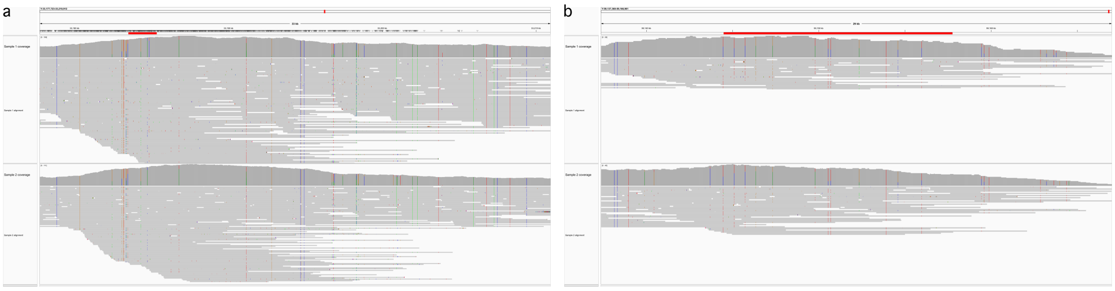

*Supplementary Figure 3. Alignments to the hemizygous region of the male specific Y indicate increases in coverage near ampliconic genes (**a**)* HSFY2 *and (**b**)* RBMY*, which are denoted by a red bar. Variant allele frequencies do not match the expected pattern for a hemizygous region, as they reflect alignments of copy number variants over the same reference location, providing an additional approach for estimating copy number.*

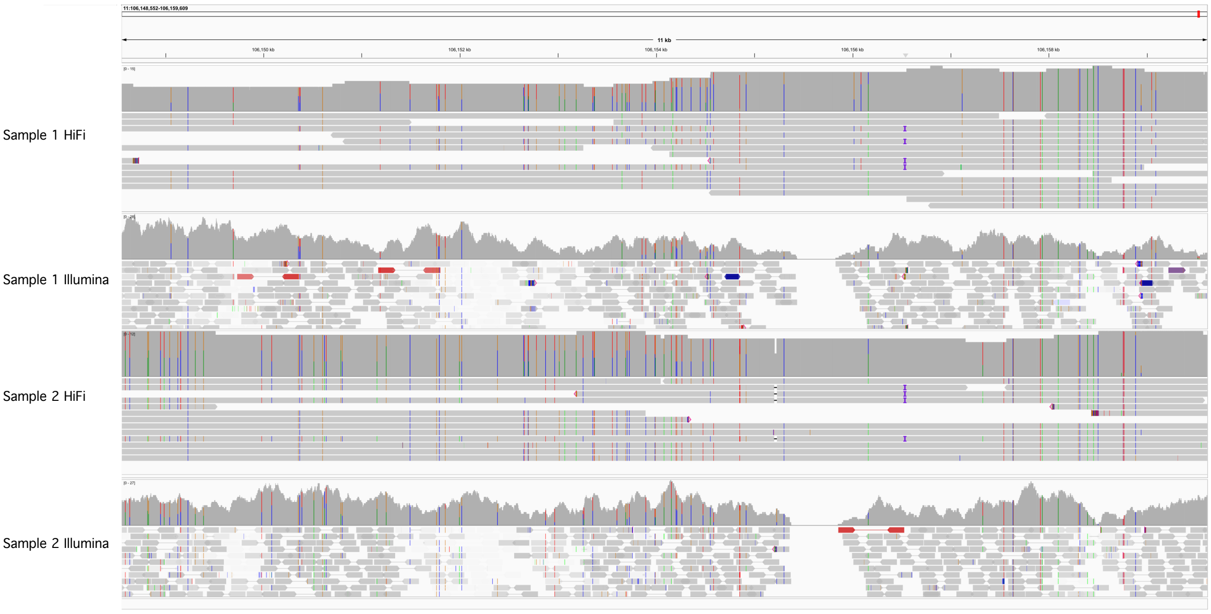


*Supplementary Figure 4. HiFi reads were able to span regions that Illumina reads had low mapping quality or other alignment issues. Signals for these variants were identified in the Illumina reads but were not called due to the alignment issues.*

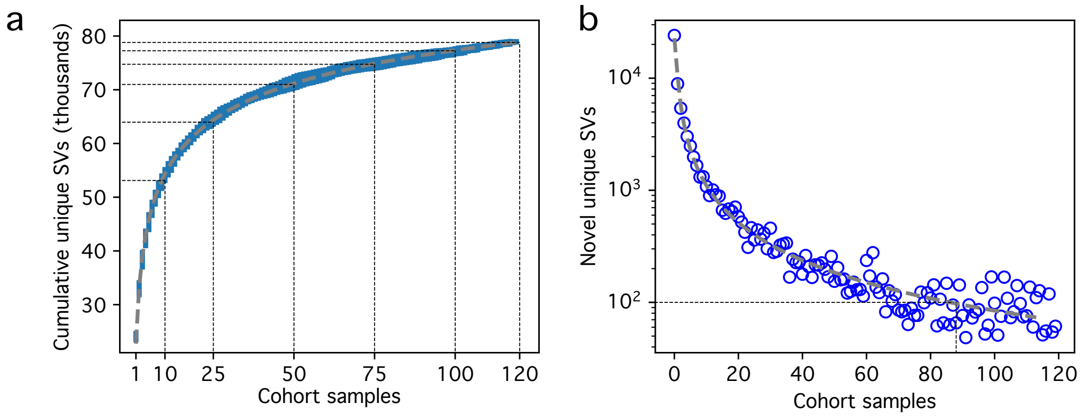


*Supplementary Figure 5. Structural variant saturation curves. (**a**) The total number of unique SVs forms a stable curve after 1,000 random shuffles of sample order. (**b**) The number of novel unique SVs rapidly decays, with fewer than 100 new SVs being found after approximately 88 samples have been included.*

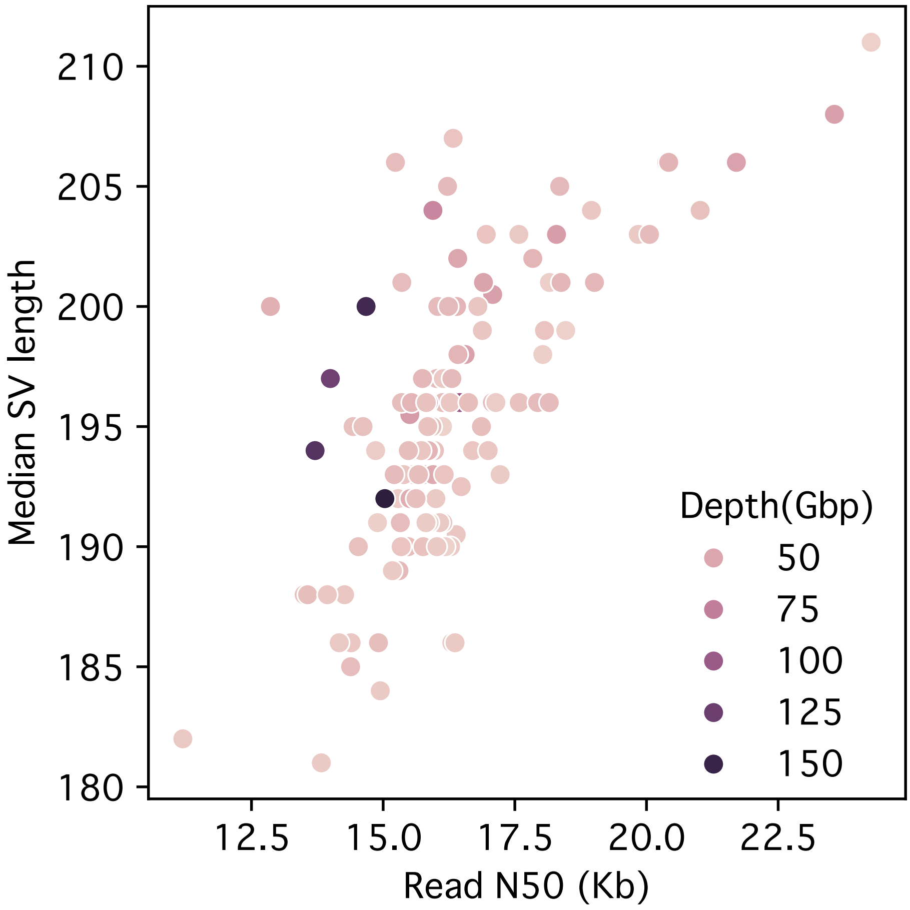


*Supplementary Figure 6. Read N50 (as well as read length) are positively correlated (Pearson's r=0.70, p=5.34e-19) with the median size of the discovered SVs, indicating long SVs are more frequently missed in samples with shorter read N50s. Rare, long SVs, only occurring in samples with low read N50, may be lost entirely.*

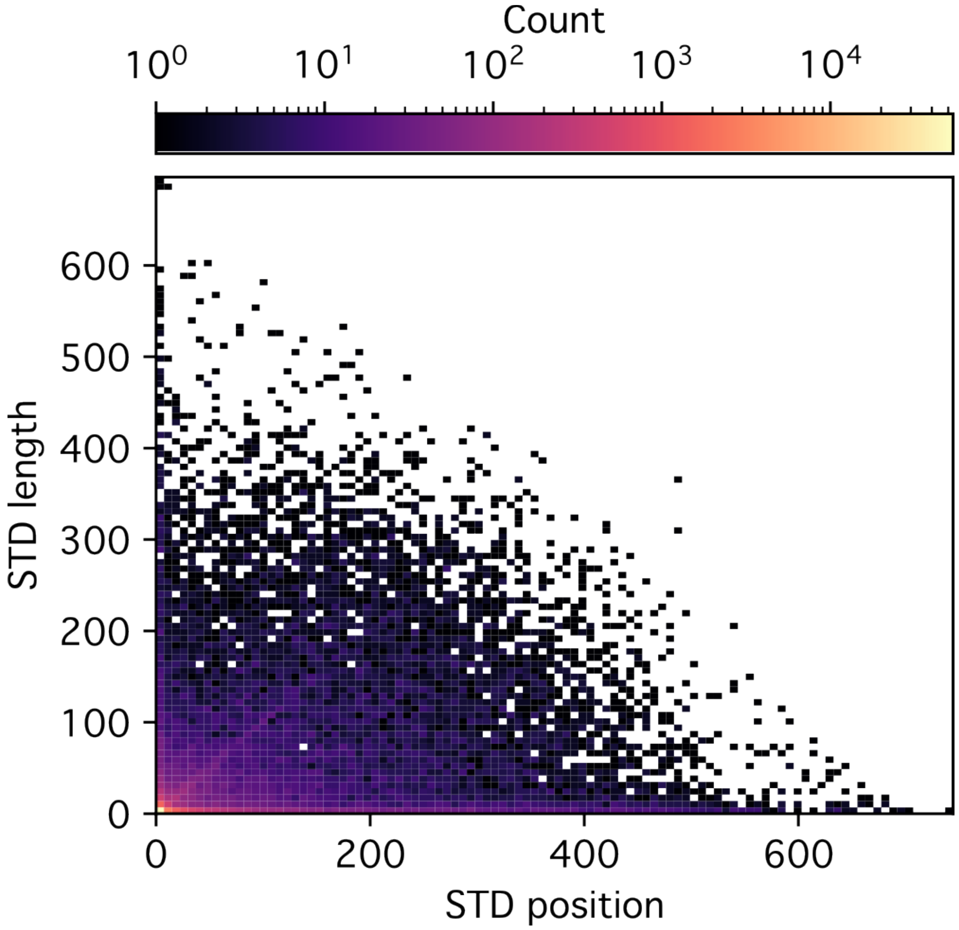


*Supplementary Figure 7. Most SVs after joint-calling had low standard deviations (STD) for both the allele start position and the allele length for the merged SVs. SVs with larger standard deviations in position and length potentially indicate merging over tandem repeat alleles with long and short motif lengths respectively.*

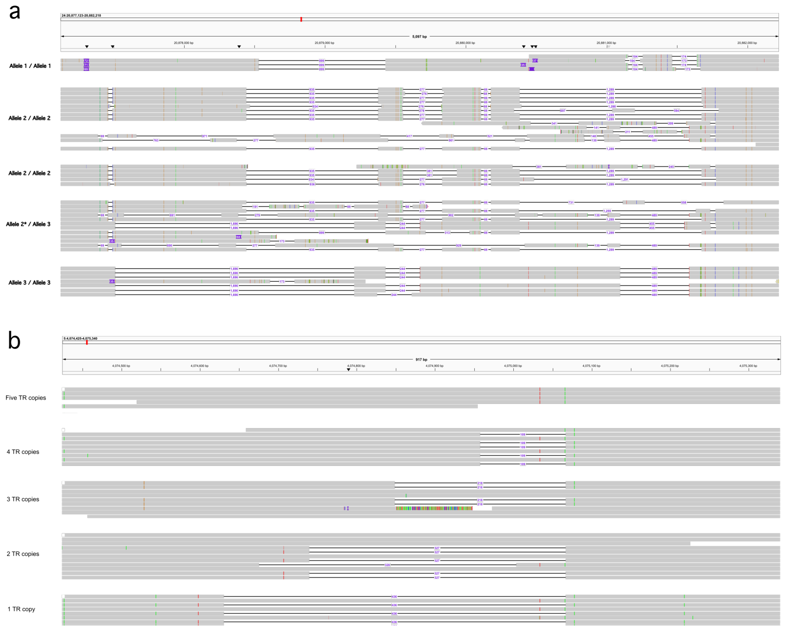


*Supplementary Figure 8. (**a**) A region annotated as a tandem repeat contained four separate SVs with three unique total deleted lengths. However, due to the different start locations of the near identical deletions exceeding the merging distance, there is a potentially "artefactual" SV which can inflate total SV numbers. (**b**) A VNTR with a motif of 109 bp, spanning five copies in the reference, was present in the cohort as a deletion covering between one and four copies. Due to merging of nearby alleles during the joint-genotyping stage, this results in a "consensus" 218 bp deletion (which was the most common allele at 39% frequency).*

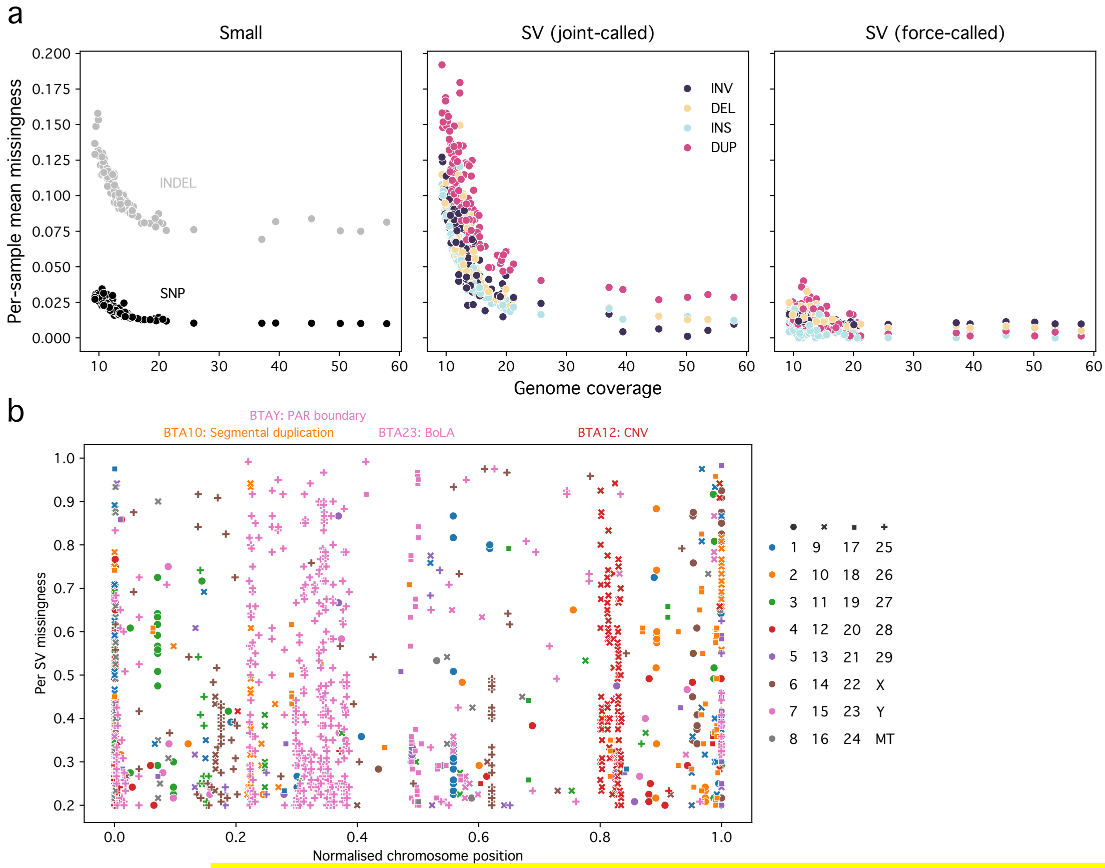


*Supplementary Figure 9. (**a**) Per-sample mean missingness is strongly correlated with low coverage for joint-called small variants and SVs, with reduced improvement after 15-20x coverage. Force-called SVs have substantially reduced missingness but still is correlated with low coverage. (**b**) Several chromosomes have SV hot spots where each SV has high missingness (e.g., BTA10, BTA12, BTA23, BTAY, etc.) corresponding to known regions of complexity/polymorphisms. Otherwise, most poorly genotyped SVs are at the start or end of chromosomes (typically centromeres or telomeres respectively), respectively at a normalised chromosome position of 0.0 or 1.0.*

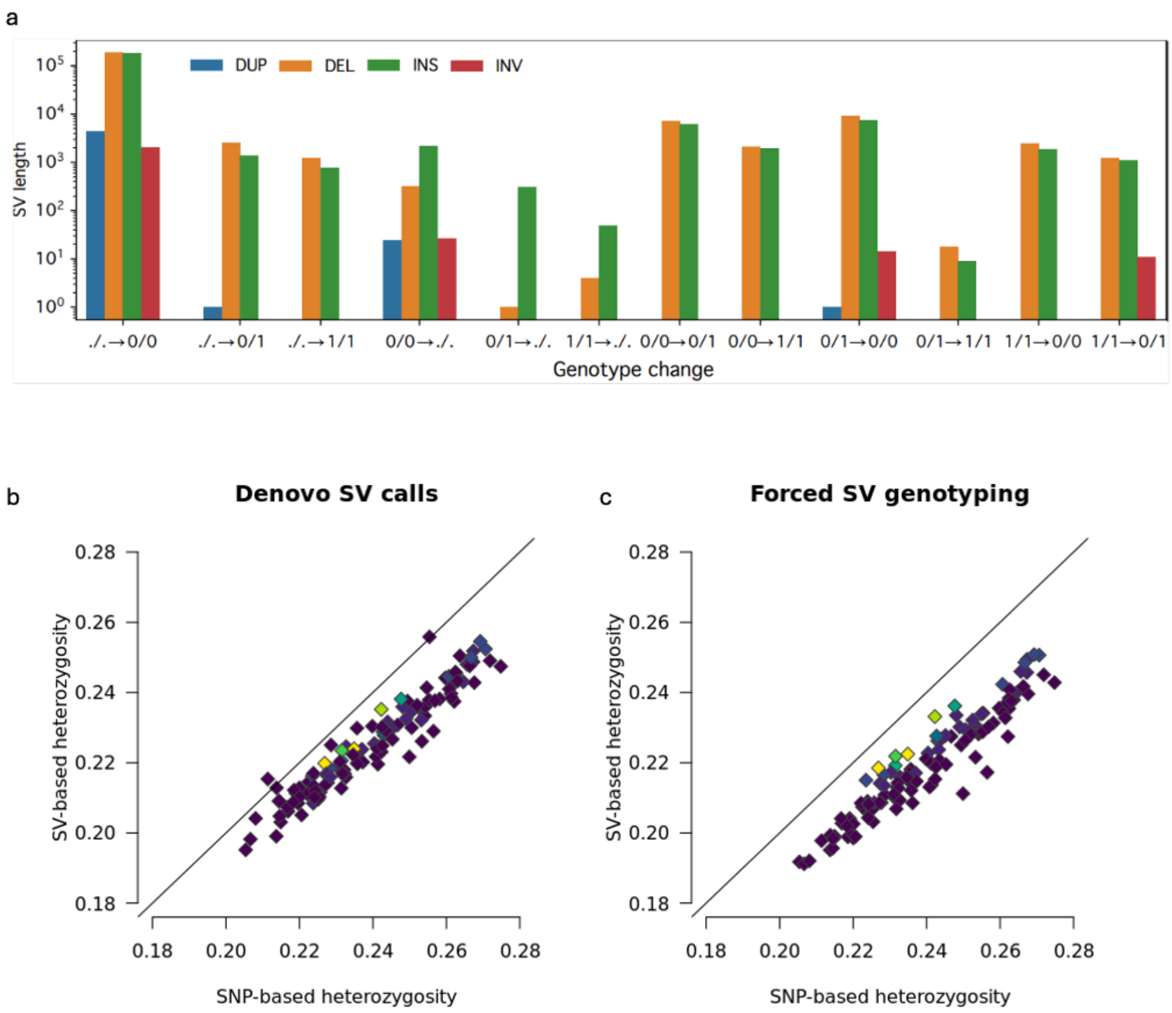


*Supplementary Figure 10. Impact of force-calling SV genotypes with sniffles2. a) Force-calling SVs with sniffles2 overwhelmingly filled in sporadically missing genotypes (note the log-scale of the y-axis). Genotype changes for insertions and deletions largely behaved similarly, except for changing existing non-reference genotypes to missing, likely occurring in regions of complex alignment. b-c) Scatterplots for the per-sample heterozygosity estimated from denovo SV calls and SNPs (b), and for the per-sample heterozygosity estimated from forced SV calls and SNPs (c). The lighter colours indicate the few samples with high coverage (>20 –fold). The fit of a linear model regressing SNP-based heterozygosity on the SV-based heterozygosity was slightly better for the denovo (adjusted R2: 0.91, p= 3.51e-64) than the forced SV (adjusted R2: 0.90, p= 2.12e-61) calls.*

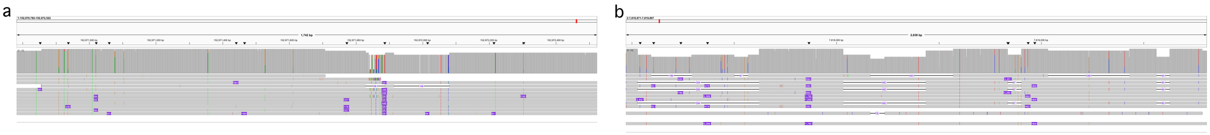


*Supplementary Figure 11. Non-missing genotype changes primarily occurred in complex regions with inconsistent alignments and clustered variation, making it challenging to manually assess the correct genotype. (**a**) IGV screenshot of a sample for which the genotype was revised from 0/0 to 1/1 for a 419 bp insertion. (**b**) IGV screenshot of a sample for which genotypes were revised for three clustered deletions: 0/1 to 0/0 (-618 bp, putatively worsened), 0/0 to 0/1 (-67 bp, putatively improved), and 0/0 (-130 bp, putatively improved).*

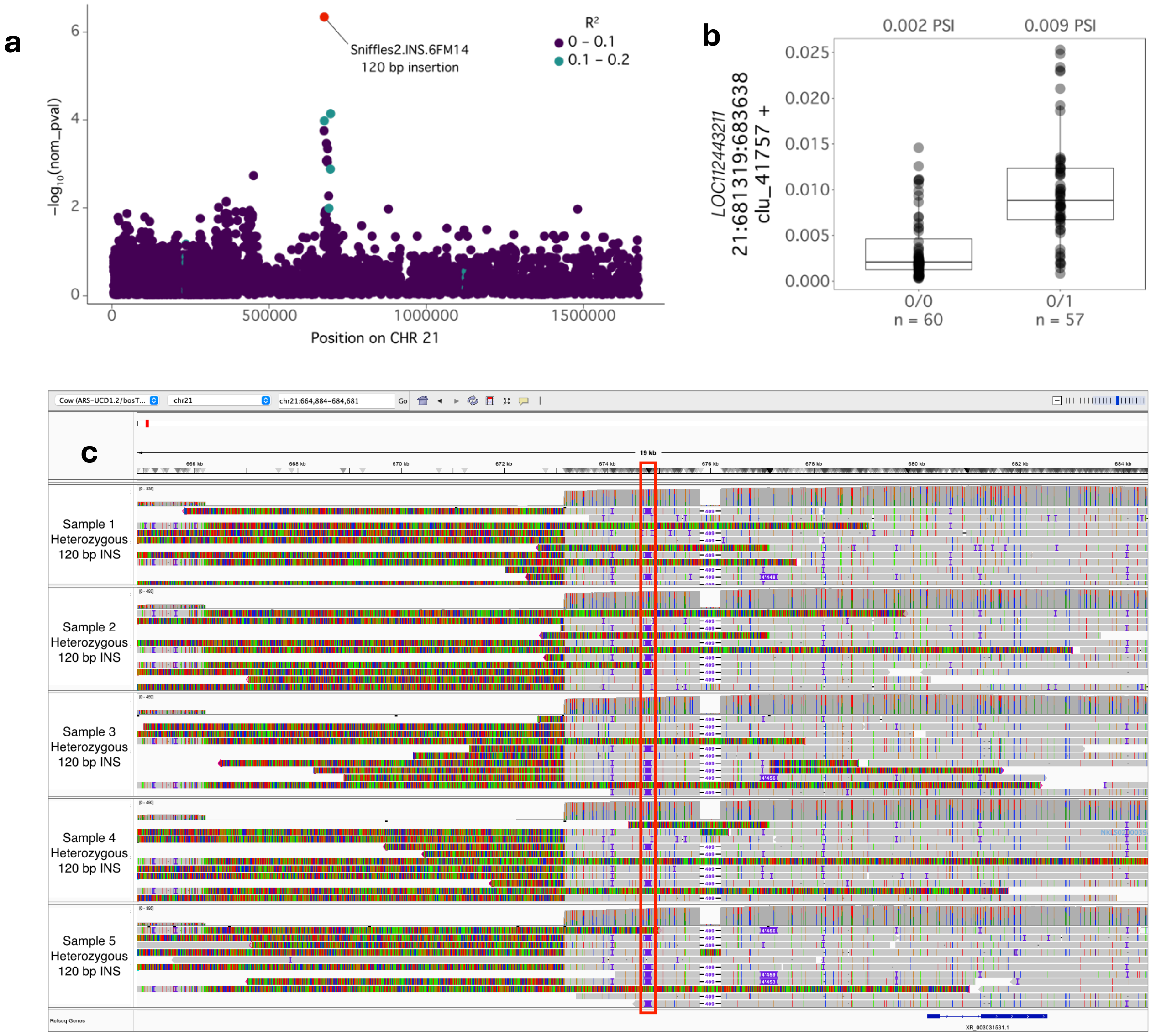


*Supplementary Figure 12. An example of a poorly tagged SV sQTL for* LOC112443211. ***(a)*** *A Manhattan plot representing the association of variants in a 1 Mb cis window with splicing variation affecting LOC112443211. The 120 bp insertion (INS; red dot) not in linkage disequilibrium ($R^2$) with any by nearby variants.* ***(b)*** *Boxplots with the genotypes for the 120 bp INS and percent spliced-in (PSI) values for the associated junction. Median PSI for each genotype is reported above, and number of samples (n) is below.* ***(c)*** *IGV alignments of 5 samples that were called heterozygous for the 120 bp INS. The INS is outlined by the red box.*

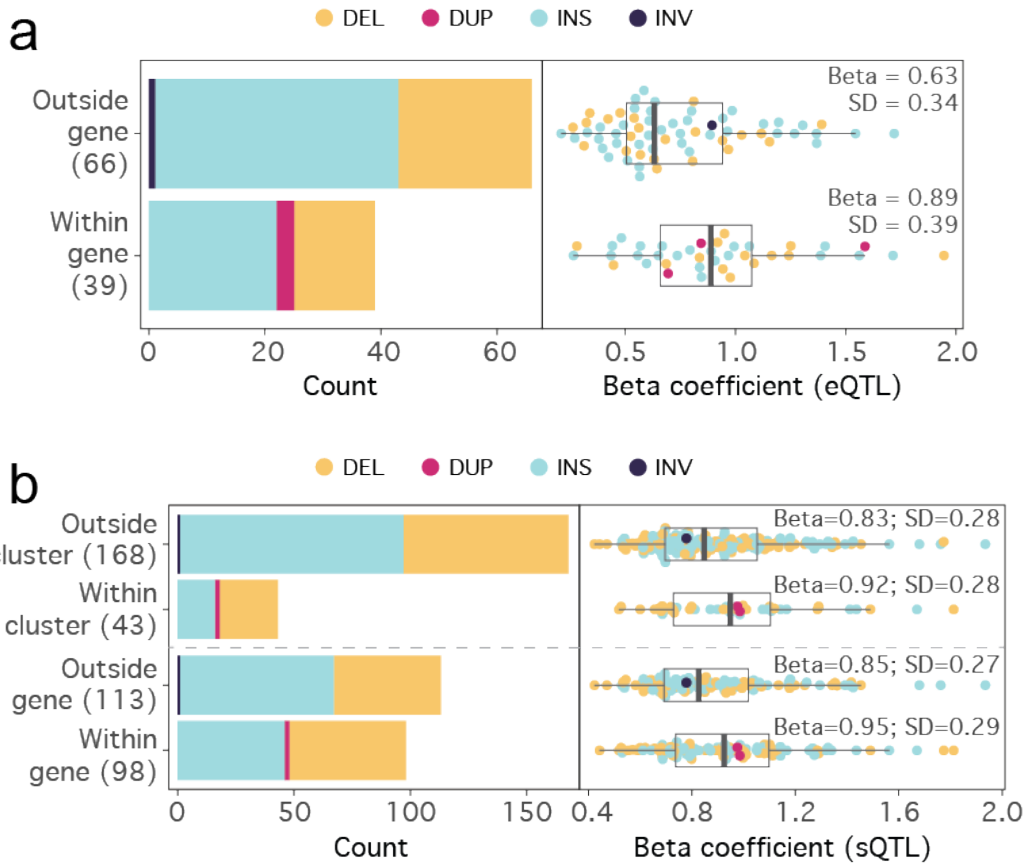


*Supplementary Figure 13. SV molQTL located within the molecular phenotype (gene body or intron cluster) have larger effect sizes, coloured by SV type (DEL – deletion, DUP – duplication, INS – insertion, INV – inversion). The left panel shows the number of SV eQTL (**a**) and SV sQTL (**b**) identified within and outside of the gene of intron cluster's body, and the right panel shows the magnitude of the effect size with the median (Beta) and standard deviation (SD) listed above.*

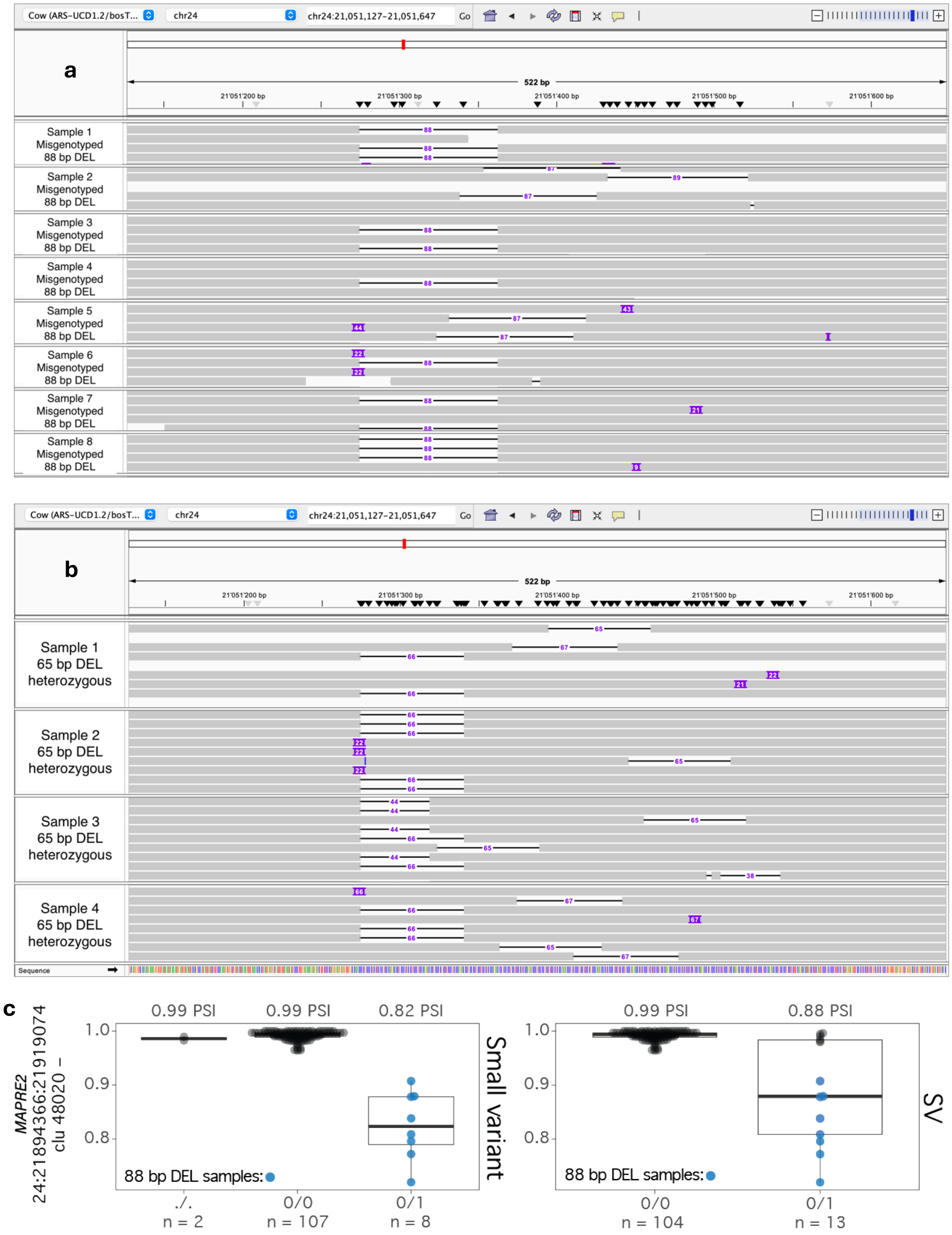


*Supplemental Figure 14. A tandem repeat (TR) that was missed as an SV sQTL because of variant merging. We identified a 22 bp tandemly repeated motif that was variable across the population. "Deletions" of different lengths (i.e. 66 bp, 88 bp, etc.) of this motif were merged into a single 65 bp deletion (DEL; hereafter referred to as the "merged SV") by sniffles2. (**a**) IGV alignments of the eight samples that were heterozygous for the 88 bp DEL. These samples were genotyped as heterozygous for the "merged SV". An example of the variability in motif length across the population is demonstrated in the IGV alignments provided in (**b**), which shows the read alignments for four samples that were heterozygous for the "merged SV". (**c**) Boxplots demonstrating the effect*

*of the top small variant (left) and the merged SV (right) genotypes on the percent spliced in (PSI) for a junction within* MAPRE2. *In both plots, the eight sample that possessed the 88 bp deletion are coloured blue. Median PSI for each genotype is reported above and sample size is reported below. The 88 bp variant was in perfect linkage disequilibrium (LD) with the small variant, while the "merged SV" was not in LD. This caused the 88 bp deletion to be missed as a potential SV sQTL and highlights the downside of merging TRs that have different functional consequences.*

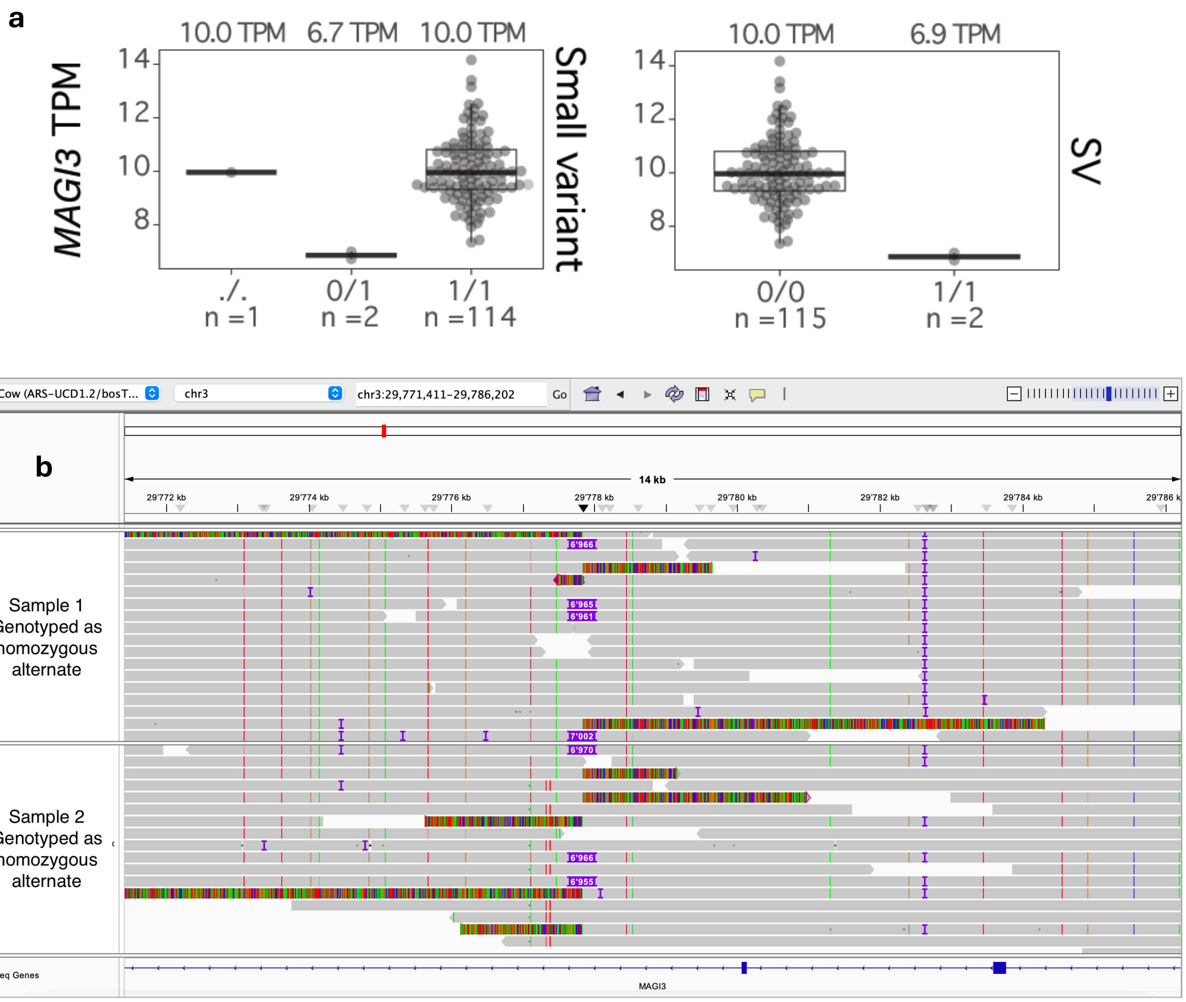


*Supplementary Figure 15. A large effect eQTL for* MAGI3 *is likely an SV. (**a**) Expression (TPM) of the top small variant for a* MAGI3 *eQTL and a 6.9 Kb insertion that was within the gene body. The small variant was prioritized as the top variant but it contained a sample with a missing genotype, whereas the SV had non-missing genotypes for all samples. Median TPM for each genotype is reported above and sample size is below. (**b**) Alignments for the two samples that were genotyped as homozygous for the 6.9 Kb insertion. The alignments indicate that both samples are misgenotyped as homozygous alternate, despite possessing numerous reads matching the reference haplotype, indicating that both samples should be genotyped as heterozygous. With the manual corrections, the top small variant is in perfect linkage disequilibrium with the SV.*

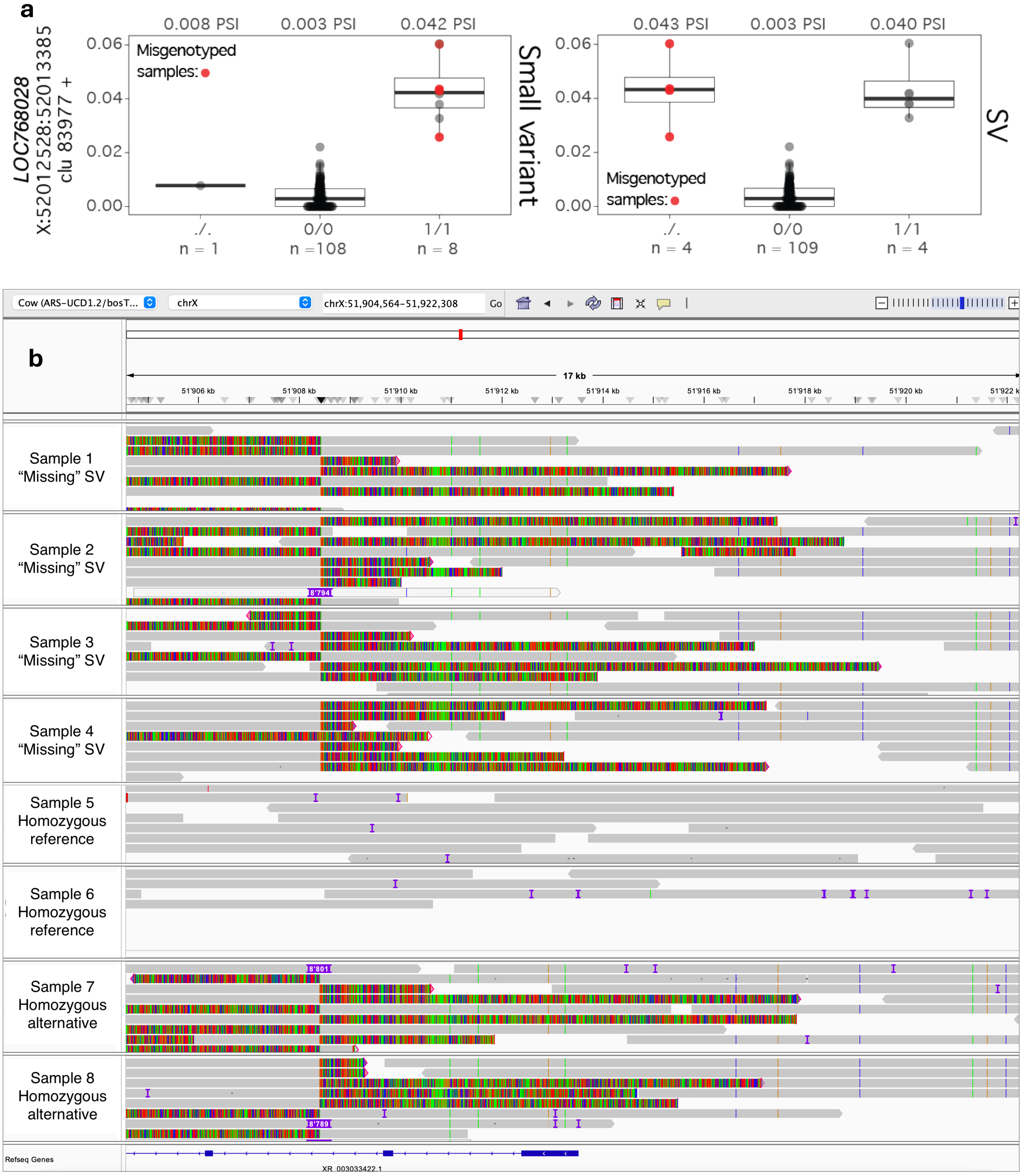


*Supplementary Figure 16. Four samples were misgenotyped as "missing" for an 8.7 Kb insertion that was associated with splicing of* LOC768028. ***(a)*** *Percent spliced in (PSI) plots for the genotypes of the top small variant of a LOC768028 sQTL, and an 8.7 Kb insertion with four samples genotyped as "missing". The samples with the "missing" SV are coloured red. Median PSI for each genotype is reported above and sample size is below.* ***(b)*** *IGV alignments for the 8.7 Kb insertion. Samples 1-4 were called as "missing", despite containing clipped reads at the location of the insertion, suggesting that these samples possess at least one haplotype with the insertion. Samples 5 and 6 are examples of correctly called homozygous reference samples, while Samples 7 and 8 are examples of correctly called homozygous alternate individuals.*

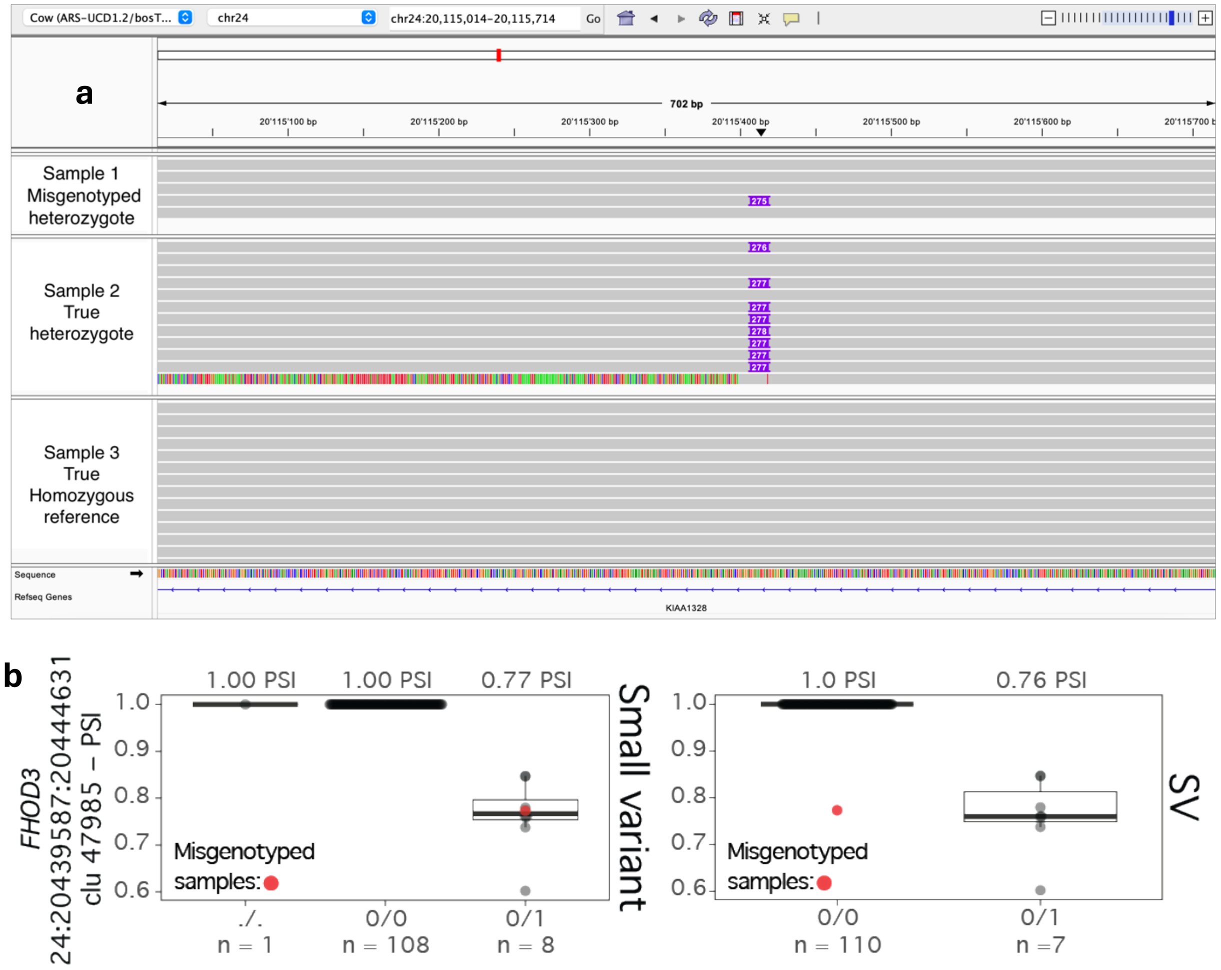


*Supplementary Figure 17. In one sample, a 277 bp insertion (INS) was misgenotyped as homozygous reference but should be heterozygous. (**a**) IGV visualization of the alignments from the individual that was misgenotyped as homozygous reference, but should be heterozygous (Sample 1), compared to a correctly called heterozygous sample (Sample 2) and a correctly called homozygous reference sample (Sample 3). (**b**) Boxplot of the 277 bp genotypes and a junction of* FHOD3*. Genotype distribution of the top small variant for the* FHOD3 *junction is on the left. The SV is in perfect linkage disequilibrium with this variant when the misgenotyped sample is corrected. Median percent spliced in (PSI) for each genotype category is reported above, and number of samples belonging to that category is listed below. The individual with the misgenotyped SV–Sample 1–is coloured red in both boxplots.*

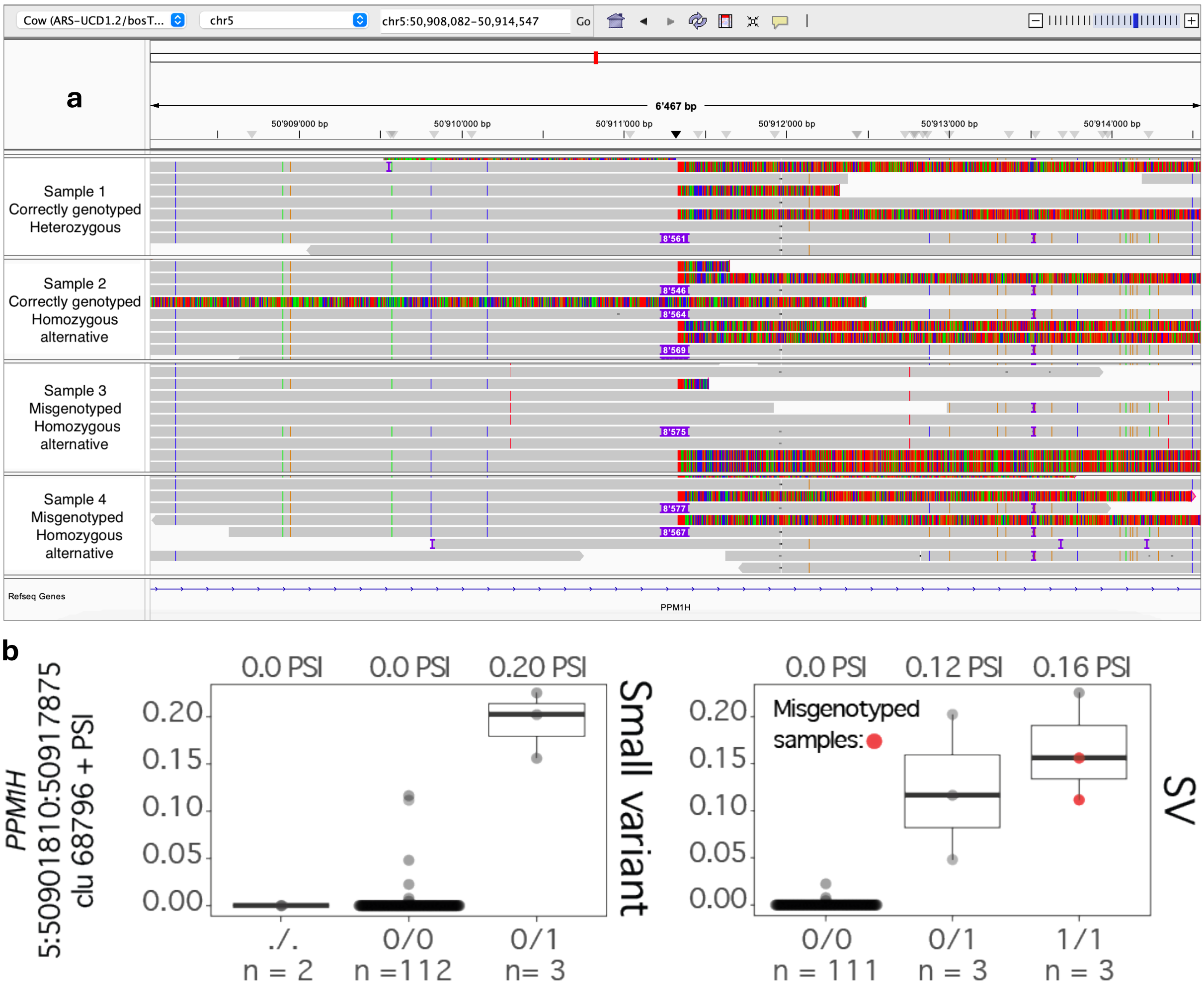


*Supplementary Figure 18. (**a**) IGV alignments for an 8,573 bp insertion. The top sample is a true heterozygous and the second sample is a true homozygous for the insertion. The bottom two samples were incorrectly genotyped as homozygous alternative, despite possessing reads that do not contain the variant. (**b**) Boxplots of the percent spliced in (PSI) for an intron cluster of* PPM1H *for the top small variant and SV with misgenotyped samples.*

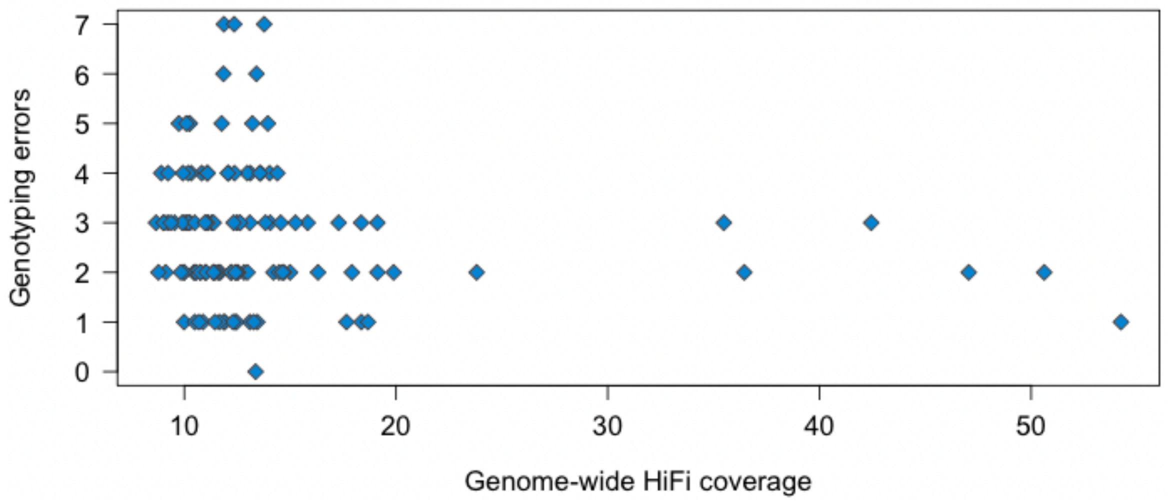


*Supplementary Figure 19. No correlation (Pearson's: r = -0.15, p = 0.09; Spearman's: rho = -0.16, p = 0.08) between the number of genotyping errors for duplications and average HiFi coverage.*

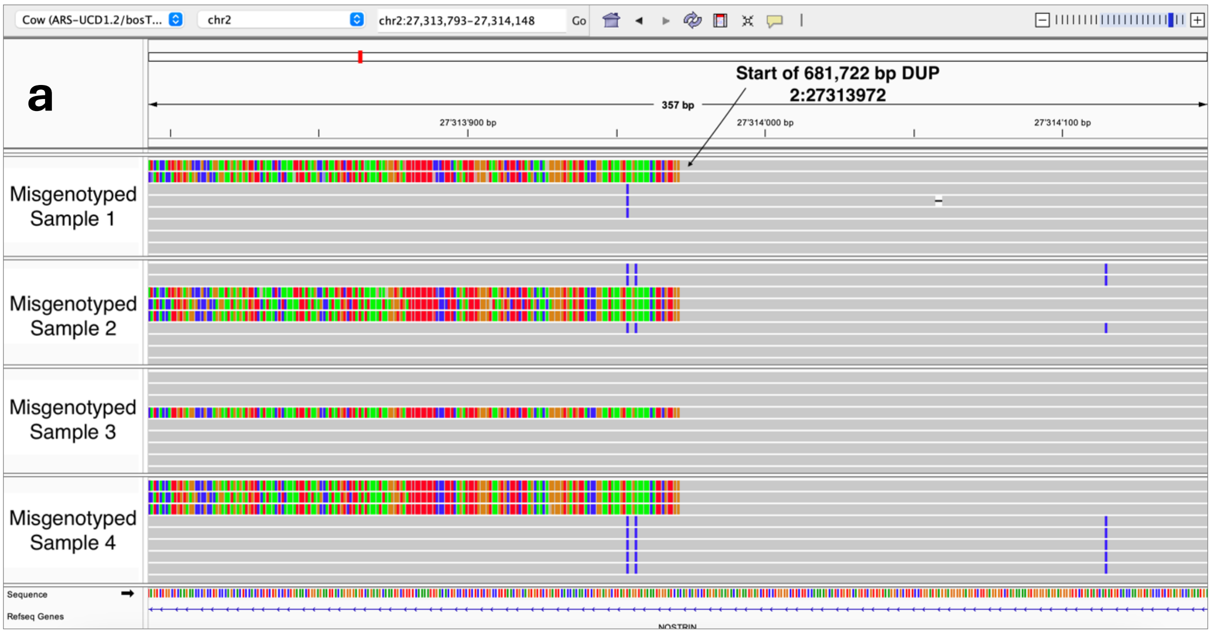


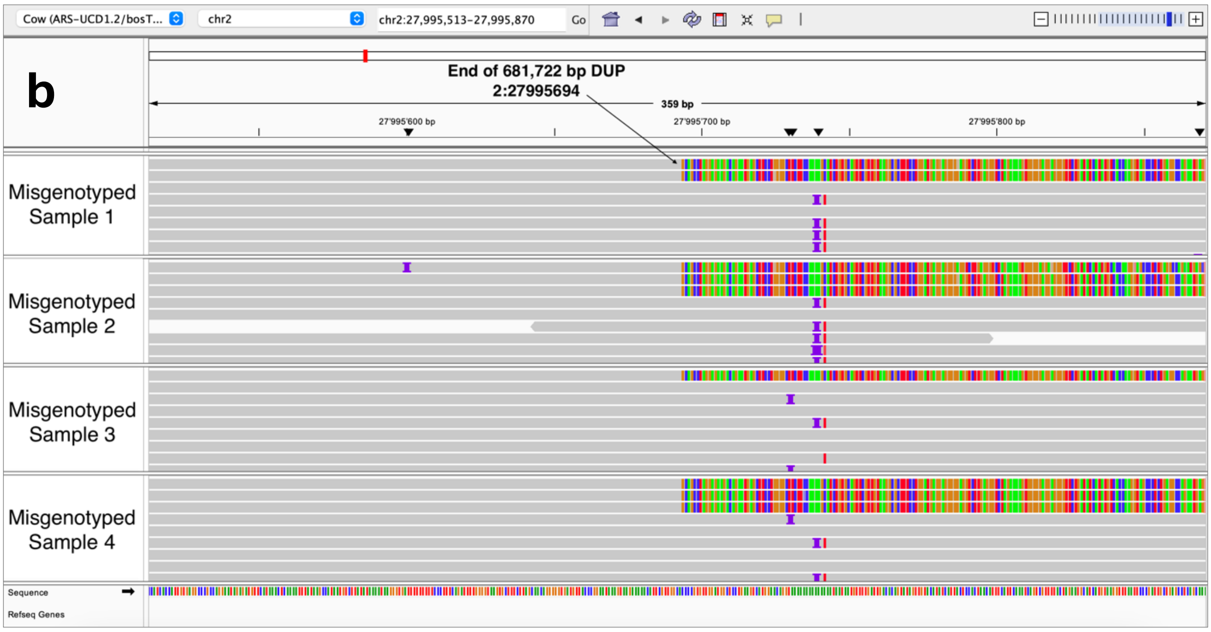


*Supplementary Figure 20. IGV alignments for four samples that show breakpoints at the beginning and end of a 681,722 bp duplication (DUP) on chromosome 2, with (**a**) showing the beginning of the duplication and (**b**) showing the end. These samples were misgenotyped as homozygous reference by sniffles2 but genotyping by coverage suggests that these four individuals are heterozygous for the deletion.*
